# MAPK ERK5 Is a Novel Regulator of MHC-I in Cancer Cells

**DOI:** 10.64898/2026.02.10.705047

**Authors:** Maria Viñas-Casas, Jose M Lizcano

**Affiliations:** Departament de Bioquímica i Biologia Molecular and Institut de Neurociències, Facultat de Medicina. Universitat Autònoma de Barcelona (UAB), Barcelona, Spain; Protein Kinases in Cancer Research Laboratory. Vall Hebron Institut de Recerca (VHIR), Barcelona, Spain

**Author notes:** **Corresponding author and address:** Jose M Lizcano. Protein Kinases in Cancer Research Laboratory, Vall d’Hebron Research Institute (VHIR), Edifici Collserola, Laboratory 202. Passeig de la Vall d’Hebron, 119-129. 08035 Barcelona, SPAIN.

**Keywords:** Map kinase, immune evasion, CD8+ T cell cytotoxicity, oncogenic signaling pathways

## Abstract

Downregulation of major histocompatibility complex class I (MHC-I) molecules is a frequent mechanism of tumor immune evasion, impairing recognition and elimination of cancer cells by cytotoxic CD8^+^ T lymphocytes. This phenotype, associated with poor prognosis and resistance to immune checkpoint blockade, is often driven by oncogenic pathways that reversibly suppress MHC-I. The MAPK ERK5 and its only upstream activating kinase MEK5 configure a unique intracellular signaling that regulates cell proliferation, differentiation and survival, and it has emerged as an oncogenic driver of different tumors. In this work, we investigated whether the ERK5 pathway contributes to cellular immunity. We used a panel of human cancer cell lines representing three well-established oncogenic contexts linked to reversible MHC-I downregulation (MYCN amplification, PTEN/PIK3CA mutations, and androgen receptor signaling), together with control cellular models exhibiting constitutively high MHC-I expression. We found that MEK5 or ERK5 pharmacologic inhibition or ERK5 targeted degradation increased MHC-I surface and total expression in low–MHC-I cells, without affecting PD-L1 levels. Conversely, ERK5 overexpression impaired MHC-I levels. Moreover, systemic administration of an ERK5 inhibitor also enhanced MHC-I expression in tumor xenografts. Mechanistically, RT-qPCR analysis showed that ERK5 or MEK5 inhibition did not significantly modify transcription of classical HLA-I genes or antigen-processing machinery components, and RNA-seq analysis did not render enrichment of MHC-I transcriptional programs in response to ERK5 inhibition. In contrast, trafficking experiments implicated the ERK5 pathway in the regulation of MHC-I lysosomal degradation, suggesting that ERK5 controls surface MHC-I through post-translational mechanisms. Notably, functional assays were carried out in co-cultures of cancer cells with tumor-specific human CD8^+^ T cells, where ERK5 inhibition sensitized MYCN-amplified or PTEN/PI3KCA-mutated cancer cells to CD8+ T cell–mediated apoptosis. These results identify ERK5 as a novel regulator of MHC-I expression in cancer cells, by regulating antigen presentation across diverse oncogenic contexts, and support a rationale for the use of ERK5 inhibitors as a strategy to improve the efficacy of immune checkpoint blockade-based immunotherapy.

## INTRODUCTION

Tumors are composed of cancer cells and a large diversity of host cells, whose interactions shape disease progression and therapy response [1]. In the tumor microenvironment, infiltrate immune cells play a critical role in controlling tumor progression, and cancer cells employ a variety of mechanisms to escape immune attack [2]. The ability of CD8+ T cells to recognize and eliminate tumor cells depends on the effective recognition of tumor antigens presented by Major Histocompatibility Complex Class I (MHC-I) molecules located on the surface of cancer cells. Therefore, many human cancers evade immune surveillance by reducing MHC-I surface expression, thereby escaping T cell recognition and cytotoxicity [3]. Loss or downregulation of MHC-I occurs through multiple mechanisms, including mutations or loss of heterozygosity of Human Leukocyte Antigen class I (HLA-I) or β2-microglobulin genes, transcriptional repression due to epigenetic silencing or impaired expression of key transcription factors, or altered trafficking and stability of MHC-I complexes at the plasma membrane, among others [4]. Clinically, MHC-I downregulation is associated with poor prognosis and represents a major mechanism of resistance to immune checkpoint blockade and other T cell–based immunotherapies [5]. Accordingly, strategies that restore or enhance MHC-I expression in tumors are being actively pursued to improve antitumor immunity [6].

The MAPK ERK5 and its only upstream activating kinase MEK5 configure a unique intracellular signaling pathway, which is activated in response to growth factors and different forms of stress, including osmotic and oxidative stress, ischemia and hypoxia (reviewed in [7, 8]). The MEK5-ERK5 axis regulates cell proliferation, differentiation and survival, and it has emerged as an oncogenic driver of different tumors [9]. Hence, overexpression of MEK5 induces tumorigenesis in murine models of lung adenocarcinoma [10], sarcoma [11] and hormone-independent breast cancer [12], whereas ERK5 elicits skin carcinogenesis [13]. Moreover, overexpression of either MEK5 or ERK5 correlates with poor prognosis and aggressiveness of numerous cancers, including prostate [14], breast [15], endometrial [16], osteosarcoma [17], and hepatocarcinoma [18], among others. Consequently, ERK5 has been proposed as a new target for anticancer targeted therapies, and ERK5 inhibitors have shown to impair the growth of several tumor xenograft mice models both, alone or in combination with chemotherapics [19].

Several oncogenic pathways have been reported to support tumor growth by regulating the expression of MHC-I molecules and associated antigen presentation components. Among these, the ERK1/2 and p38 MAP kinase pathways have been implicated in tumor immune evasion, in part by regulating MHC-I surface expression through transcriptional mechanisms in several models of solid cancers [20, 21]. In this study, we investigated the contribution of oncogenic ERK5 signaling to regulation of MHC-I expression in cancer, using a panel of cancer cells harbouring well-established key oncogenic alterations associated with reversible MHC-I downregulation. We show that pharmacologic inhibition or targeted degradation of ERK5 increases MHC-I surface expression in cancer cells with low basal levels, without affecting PD-L1 expression, while ERK5 overexpression impairs MHC-I levels. Relevantly, ERK5 inhibition sensitized tumor cells to CD8+ T cell–mediated cytotoxicity and restored MHC-I expression across diverse oncogenic context, such as PI3KCA/PTEN mutations or NMYC amplification. Our findings identify ERK5 as a novel regulator of MHC-I and highlight its potential as a therapeutic target to enhance antitumor immunity.

## Methods

### Inhibitors, targeted degraders and other reagents

ERK5 inhibitor JWG-071 (MedChemExpress), MEK5 inhibitor GW284543 (CliniSciences), ERK5 proteolysis-targeting chimera (PROTAC) INY-06-061 and its negative control INY-06-089 (a gift from Prof. N. Gray, [22]) and proteasome inhibitor MG-132 (MedChemExpress) were resuspended in dimethyl sulfoxide (DMSO, Sigma). Anterograde transport inhibitor brefeldin A (Sigma) was resuspended in absolute ethanol. Lysosome inhibitors chloroquine (Sigma) and bafilomycin A (Enzo) were resuspended in water. Interferon-γ (IFN-γ, PeproTech) and TNF-α (PeproTech) were diluted in H_2_O 0.1% BSA.

### Cells, cell culture and transfection

A549 (Cat#CCL-185) non-small cell lung cancer cells, IMR-32 (Cat#CCL-127) neuroblastoma cells and LNCaP (Cat#CRL-1740) prostate cancer (PCa) cells were purchased from ATCC. Ishikawa (Cat#99040201) endometrial cancer (EC) cells were from ECACC and were purchased from Sigma. Ishikawa ERK5 CRISPR-KO cells were previously described [23]. ARK1 EC cells and SW620 colorectal cancer cells were a gift from Dr. E. Colas and Dr. T. Aasen (VHIR, Barcelona), respectively. ARK1 and A549 cells were maintained in Dulbecco’s Modified Eagle’s Medium (DMEM; Gibco) supplemented with 10% fetal bovine serum (FBS; Merck) and 1% Penicillin/Streptomycin (Pen/Strep; Gibco). SW620 cells were cultured in DMEM supplemented with 10% FBS and 1mM sodium pyruvate (Gibco). IMR-32 cells were cultured in Iscove′s Modified Dulbecco′s medium (IMDM; Gibco) with 20% FBS (Merck), 1% Pen/Strep (Gibco) and 1% Insulin Transferrin-Selenium (Gibco). Ishikawa cells were maintained in Modified Eagle Medium (MEM; Gibco) with 5% FBS (Merck) and 1% Pen/Strep. LNCaP cells were cultured in Roswell Park Memorial Institute Medium (RPMI-1640; Gibco) supplemented with 10% FBS (Merck), 1% Pen/Strep (Gibco), nonessential amino acids (NEAA; Sigma), 1mM sodium pyruvate and HEPES (Gibco). All cell lines were tested for mycoplasma contamination every three months. Ishikawa cells were transfected using Lipofectamine2000TM (Thermofisher) as described before [24]. The pEBG2T vector encoding the GST-tagged human ERK5 protein was previously described [25].

For 3D cultures of Ishikawa cells, a Matrigel (Corning) layer of approximately 1–2mm thick was added to each well of a 48-well plate. After solidification at 37 °C, 10,000 Ishikawa cells were seeded per well in basal medium (DMEM/F-12, with 1mM sodium pyruvate, HEPES and 1% Pen/ Strep) supplemented with 2% dextran-coated charcoal-stripped serum (Gibco) and 3% Matrigel. 3D cultures were then left to grow for 5 days, until the spheroids reached an approximate diameter of 100 μm, prior to incubation with indicated treatments, disaggregation and MHC-I staining.

### Cell lysis and immunoblotting

For immunoblot analysis, cells were lysed in ice-cold RIPA buffer supplemented with 50mM NaF, 5mM sodium-pyrophosphate and 1mM sodium-orthovanadate. Cell lysates were sonicated and centrifuged at 12.000 g for 12min at 4°C, and supernatants were stored at −20°C until use. Protein concentration was determined using Pierce™ Coomassie Plus reagent (Thermofisher). Proteins were resolved in SDS-PAGE gels and electrotransferred onto nitrocellulose membranes (Merck). After incubation with the appropriated primary antibody, detection was performed using horseradish peroxidase-conjugated secondary antibodies and enhanced chemiluminescence reagent (Bio-Rad). Primary and secondary antibodies used are given in **Supplementary table 1**. Full and uncropped western blots are provided as supplemental material.

### Tumor cell-T cell and cancer cell in vitro co-culture system

We followed the protocol described by Ma et al [26]. Briefly, IMR-32 and Ishikawa cell lines were trypsinized and resuspended in PBS to obtain single-cell suspensions (2×10^7^ cells/ml). Cell lysates were prepared by 5 cycles of repeated freeze-thaw (alternating between dry ice and 37ºC). After centrifuging at 2000 g for 10 min, the supernatant was collected and filtered through 0.22 μm filter to generate a tumor-specific antigen solution, which can be stored at −80°C. The day before lymphocytes preparation, T75 culture flasks were coated with 3ml of the tumor-specific antigen solution (20 μg/mL in PBS) and incubated overnight at 4°C. The next day, PBMCs from healthy donors were separated by Ficoll (Lymphoprep, Palex) density centrifugation. Erythrocytes were depleted using RBC lysis buffer (BD). PBMCs were then washed and resuspended in serum-free lymphocyte medium (X-vivo 10, Lonza) at a concentration of 1×10^6^ cells/ml, followed by seeded into tumor-specific antigen coated flasks for continuous incubation. In the first 48 h of incubation, 600U/ml GM-CSF (PeproTech) and 400U/ml IL-4 (PeproTech) were added to promote the differentiation of mature antigen-loaded dendritic cells (DCs). Subsequently, 1200U/ml IFN-γ (PeproTech), 1000U/ml IL-2 (PeproTech) and 150 ng/mL anti-CD3 antibody (Invitrogen) were added to stimulate T cell activation. Cells were then continuously cultured for 15 days, with fresh media containing IL-2 replaced every second day. The cell density was maintained at 1–2× 10^6^ cells/ml. At the end of the culture period, a large population of DC-CTLs was isolated using a magnetic-based beads positive selection system (CD8 microbeads, Miltenyi Biotec) to obtain CD8+ T cells. For in vitro co-culturing experiments, IMR-32 or Ishikawa cells were seeded in 6-well plates. The following day, cells were treated with ERK5i or vehicle control (DMSO) for 24 hours. CD8+ T cells were then added to the tumor cells at different E:T (Effector-Target) ratios and co-cultured overnight. Apoptosis was assessed by flow cytometry.

### Flow cytometry

For cell surface staining, human cancer cells were trypsinized into a single-cell suspension (5 minutes at 37ºC for monolayer cultures and 1 hour at 37ºC for 3D cultures). Cells were then washed with FACS buffer (PBS with 2% FBS) and stained with fluorochrome-conjugated antibodies and a Live/Dead fixable dead cell stain kit (near-IR fluorescent, Invitrogen) for 30 minutes at 4ºC in the dark. The antibodies used were obtained from BioLegend: Pacific Blue anti-human HLA-A,B,C (W6/32) and APC anti-human CD274 (B7-H1, PD-L1).

For apoptosis analysis, the Annexin-V Apoptosis detection kit (Invitrogen) was used. Briefly, the supernatant from co-cultures (containing CD8+ T and dead human cancer cells) was collected, and the remaining adherent cells were harvested by trypsinization (5 minutes at 37ºC). All cells were first washed with FACS buffer and then stained with fluorochrome-conjugated antibodies Pacific Blue anti-human HLA-A,B,C (W6/32) and PerCP anti-human CD3 (HIT3a, BioLegend) for 30 minutes at 4ºC in the dark. After staining, cells were washed twice with Annexin binding buffer and finally incubated with Annexin-V-APC and propidium iodide (PI) for 15 minutes at room temperature in the dark. Samples were acquired on a BD LSRFortessa flow cytometer (BD Biosciences) and data were analyzed using FlowJo Software (vX.0.7, TreeStar).

### RT-qPCR analysis

Total RNA was extracted from human cancer cells using RNeasy kit (Qiagen), following the manufacturer’s instructions. Total cDNA was obtained using the iScript™ cDNA Synthesis kit (Bio-Rad). The resulting cDNA was then subjected to real-time quantitative PCR assay using SYBR Green Supermix (Bio-Rad) in a Roche LightCycler system. All qPCR assays were performed in three replicates of each sample and from at least three independent experiments. RT-qPCR reaction conditions were as follows: 95°C for 5 min, 40 cycles at 95°C for 5 s, 60°C for 20 s, and 60°C for 30 s. The genes studied were *HLA-A, HLA-B, HCL-C, B2M, TAP1, TAP2, KLF2, CDKN1A* (p21) and *TBP* was used as the housekeeping gene. qPCR primer sequences are given in **Supplementary Table 2**. Relative gene expression levels were calculated using the 2^-ΔΔCt method.

### RNAseq analysis

We used the RNA sequencing datasets generated from Ishikawa endometrial cancer cells treated for 24 h with either vehicle or the ERK5 inhibitor JWG-071 ([23], GEO database repository under accession code GSE239920). New functional enrichment analyses were performed using Gene Set Enrichment Analysis (GSEA) software.

### Immunohistochemistry staining

Paraffin-embedded samples from human endometrial cancer tumor xenografts previously generated [16] were sectioned at 3 μm thickness sections and stained with haematoxylin/eosin, and with an anti-HLA CLASS1 ABC mAb (EMR8-5, Abcam). IHC staining was performed by the Histopathology Unit from the CNIO (Madrid, Spain).

### Schematics and statistical analysis

The scheme in Figure 3C, 4C and 5A were created with BioRender (BioRender.com). Individual data points are shown in the bar plots. Under otherwise stated, data are presented as the mean ± standard deviation (S.D.) of *n*=3 independent experiments. The statistical tests used are indicated in figure legends. Statistical significance was determined using two-tailed Student’s *t*-test, Mann-Whitney test, one-way ANOVA followed by Bonferroni multiple comparisons test or two-way ANOVA followed by Tukey multiple comparisons test, and was set at *p*<0.05. All analyses were performed using GraphPad Prism 8 software.

## Results

### MEK5-ERK5 signaling pathway modulates surface MHC-I expression in human cancer cell lines that exhibit downregulated MHC-I membrane levels

To investigate the role of ERK5 in cancer cell immune evasion mechanisms, we used flow cytometry analysis to assess the membrane levels of the Major Histocompatibility Complex Class I (MHC-I) across six human cancer cell lines: IMR-32 (neuroblastoma, NBL, *MYCN* amplified), Ishikawa (endometrioid cancer, EC, PTEN-null), ARK1 (serous EC. PI3KCA mutant), LNCaP (androgen-responsive prostate cancer, PTEN-mutant), A549 (non-small cell lung cancer, *KRAS*-mutant) and SW620 (colorectal cancer, *KRAS*-mutant) (**Supplementary Table 3**).

Among the cell lines studied, IMR-32, Ishikawa and LNCaP cells exhibited downregulated (low basal) MHC-I surface expression. In these cells, ERK5 inhibitor JWG-071 (ERK5i, [27]) or MEK5 inhibitor GW284543 (MEK5i, [28] increased the percentage of cells expressing MHC-I as well as the MHC-I mean fluorescence intensity (MFI) (**Fig. 1A**). In contrast, ARK1, A549 and SW620 cells showed high basal MHC-I surface expression, with all cells being positive for MHC-I staining. In these cells, ERK5 or MEK5 inhibition did not alter the MHC-I MFI values (**Fig. 1B**). PD-L1 upregulation is a well-known mechanism for the immune evasion of cancer cells through impairment of the cytotoxic activity of CD8+ T cells [2]. Hence, we also assessed PD-L1 membrane expression in the low MHC-I-expressing cells (IMR-32, Ishikawa and LNCaP). These cells exhibited increased PD-L1 expression in response to the canonical inducers INF-γ, TNF-α or the proteasomal inhibitor MG-132 (**Fig. 1C**). However, ERK5 or MEK5 inhibition did not alter either the percentage of PD-L1+ cells or the PD-L1 MFI values (**Fig. 1C**).

**Figure 1.**
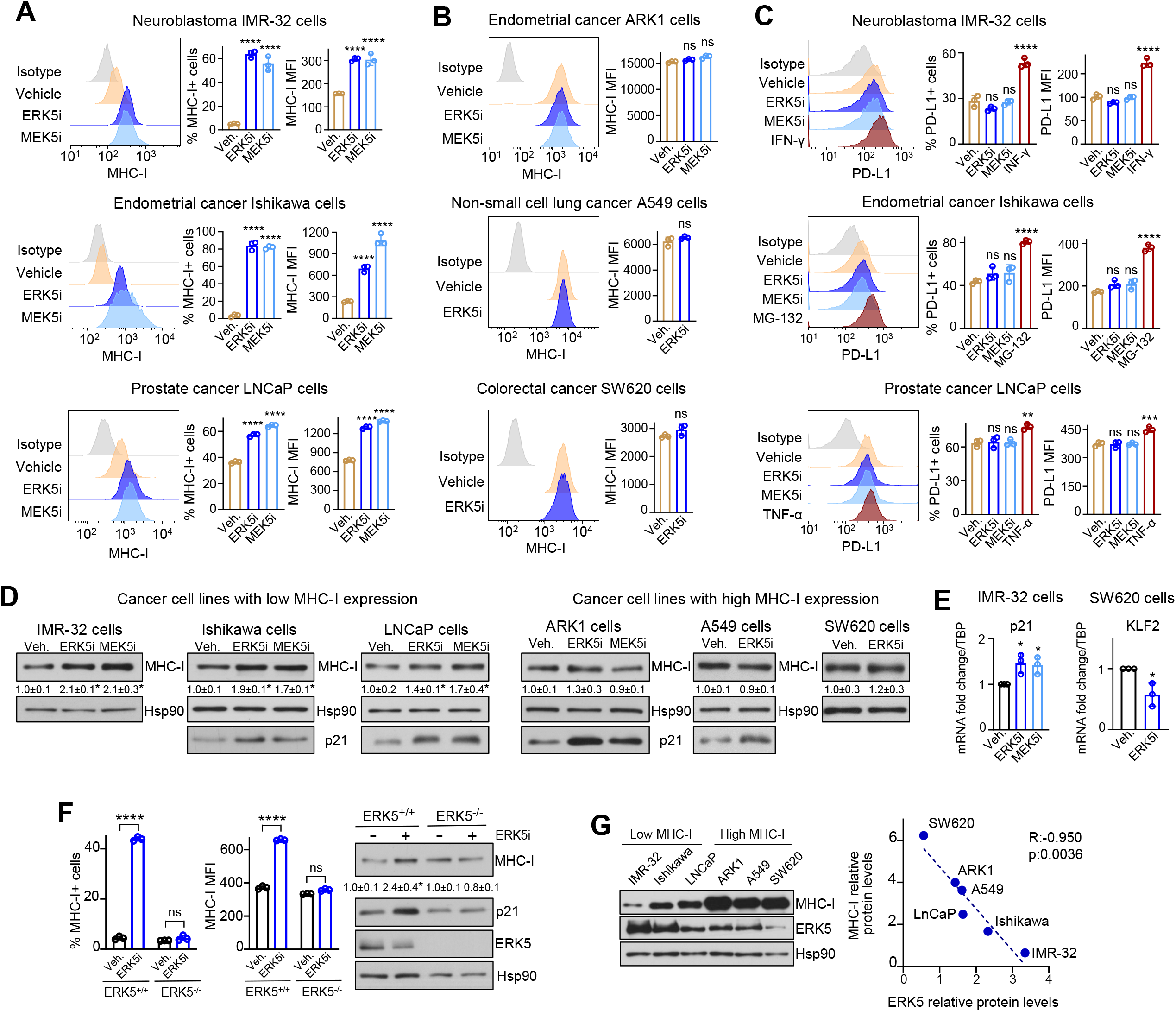
ERK5 and MEK5 inhibitors enhance MHC-I surface expression in cancer cells without affecting PD-L1. (**A**) Effect of ERK5/MEK5 inhibitors in cancer cells with low basal MHC-I expression. IMR-32, Ishikawa and LNCaP cells were treated for 24h with vehicle, 5μM JWG-071 (ERK5i) or 5μM GW284543 (MEK5i), and MHC-I surface expression was assessed by flow cytometry (FC). Left panels show representative FC histograms, and right panels show percentages of cells expressing MHC-I and the mean fluorescence intensity (MFI) values. (**B**) Effect of ERK5/MEK5 inhibitors in cancer cells with high basal MHC-I expression. ARK1, A549 and SW620 cells were treated as in (**A)**, and MHC-I membrane expression was determined by FC. Representative FC histograms and MFI values are presented. (**C**) Effect of ERK5/MEK5 inhibitors in membrane PD-L1 expression in cancer cells with low basal MHC-I levels. IMR-32, Ishikawa and LNCaP cells were treated as in (**A)**, or with 20ng/mL IFN-γ or 8μM MG-132 or 10ng/mL TNF-α as positive controls. PD-L1 surface expression was measured by FC. Left panels show representative FC histograms, and right panels show percentages of cells expressing PD-L1 and MFI values. (**D**) Effect of ERK5/MEK5 inhibitors on total MHC-I protein levels. The indicated cells were treated with vehicle, 5μM JWG-071 or 5μM GW284543 for 24h, and MHC-I protein levels were determined by immunoblot. The numbers indicate relative MHC-I protein levels referred to vehicle (mean ± SD from three independent experiments). Immunoblot (**D**) or qPCR analysis of p21 mRNA levels, as well as relative *KLF2* gene expression (**E**) were used as controls for ERK5 inhibition. mRNA levels were normalized to *TBP* mRNA levels. (**F**) Effect of 5μM ERK5i on MHC-I expression in Ishikawa ERK5 wild type (ERK5^+/+^) and ERK5 CRISPR-KO (ERK5^-/-^) cells. Cells were treated 24h and MHC-I membrane levels were assessed by FC (left panels show percentages of cells expressing MHC-I and MFI values), and total MHC-I protein levels by immunoblot (right panels). p21 levels were used as a control for ERK5 inhibition (**G**) Total MHC-I and ERK5 protein levels inversely correlate in cancer cells. Immunoblot analysis show MHC-I and ERK5 expression in the indicated cells. Right plot shows Pearson’s pairwise correlation analysis of relative levels of ERK5 and MHC-I protein expression in the cell lines tested. Data represents mean ± SD from three independent experiments. ns, not significant; * *p* < 0.05; ** *p* < 0.005; *** *p* < 0.001; **** *p* < 0.0001, as determined by one-way ANOVA followed by Bonferroni multiple comparisons test.

We also investigated the impact of ERK5 pathway inhibitors on total MHC-I protein levels by immunoblot analysis. Consistent with the flow cytometry data, both MEK5 and ERK5 inhibitors increased MHC-I protein levels in low basal MHC-I-expressing cells, but not in cells with high basal MHC-I expression (**Fig. 1D**). As controls, both ERK5i and MEK5i induced the expression of the established downstream target p21 (**Fig. 1D-E)**, or reduced the expression of the transcriptional target *KLF2* (**Fig. 1E**). Moreover, the ERK5 inhibitor JWG-071 did not alter the percentage of MHC-I+ cells or MFI values in Ishikawa ERK5 CRISPR-KO cells, reflecting the specificity of this compound (**Fig. 1F**). Regarding the effect of ERK5 pathway inhibition on known regulators of MHC-expression, ERK5 and MEK5 inhibitors did not alter the phosphorylation status of ERK1/2 or Akt proteins, but affected c-Myc protein levels in all cell lines tested except IMR-32 cells (**Supplementary Fig. 1)**. Of relevance, immunoblot analysis across the cancer cell line panel revealed a striking inverse correlation between MHC-I and ERK5 protein levels (Pearson R:-0.950, *p:*0.0036; **Fig. 1G**), further supporting a role for ERK5 in regulating MHC-I expression. A similar analysis for phosphorylated ERK5 could not be performed, as the phospho-specific antibody did not yield a detectable signal under our experimental conditions (**Supplementary Fig. 2**).

Together, these findings show that inhibitors of the MEK5-ERK5 pathway increases MHC-I membrane levels in cancer cells with downregulated MHC-I, without affecting PD-L1 surface expression.

The effect of ERK5 pathway inhibitors on MHC-I expression in cancer cells was further validated by either abolishing or overexpressing the ERK5 protein. We used the PROTAC degrader INY-06-061, which specifically targets ERK5 for ubiquitylation and proteasomal degradation [22]. As negative control PROTAC, we used the INY-06-089 compound, which shares the same structure as the functional compound but lacks the ability to bind ERK5. As expected, the ERK5 degrader, but not the negative control compound, completely abolished ERK5 protein expression in the IMR-32, Ishikawa or LNCaP cancer cells (**Fig. 2A-C**), without affecting the total and phosphorylated levels of relevant proteins such as ERK1/2 or Akt (**Supplementary Fig. 3**). More importantly, INY-06-061 significantly increased the percentage of cells expressing membrane MHC-I and the MHC-I MFI values, as well as of total MHC-I protein levels, in the three low basal MHC-I-expressing cell lines (**Fig. 2A-C)**. As observed for the ERK5/MEK5 inhibitors, the ERK5 PROTAC had no effect on the membrane and total MHC-I levels in cells with high low basal MHC-I expression (**Fig, 2D**). Conversely, overexpression of ERK5 protein in Ishikawa EC cells resulted in lower percentage of MHC-I+ cells, MHC-I MFI value and of total MHC-I protein levels (**Fig. 2E**), suggesting an immunosuppressive role for ERK5 in suppressing MHC-I membrane expression, at least in EC cells.

**Figure 2.**
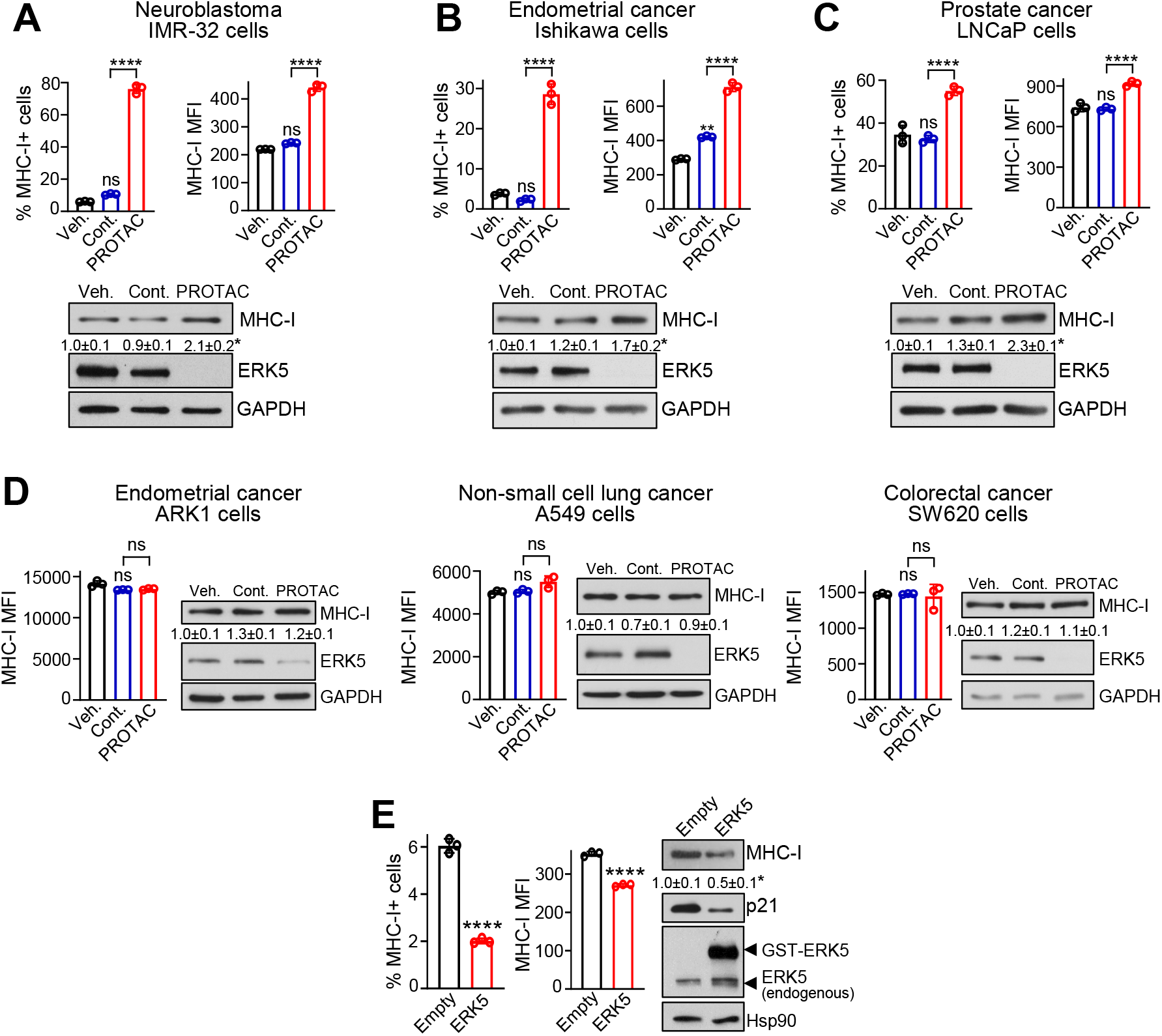
ERK5 targeted degradation upregulates MHC-I membrane levels in cancer cells, whereas ERK5 overexpression reduces MHC-I expression. (**A-C**) ERK5 PROTAC degrader enhances MHC-I membrane levels in cancer cells with low basal MHC-I surface expression. IMR-32 (**A**), Ishikawa (**B**) and LNCaP (**C**) cells were treated for 24 h with vehicle, 300nM PROTAC negative control INY-06-089, or 300nM ERK5 PROTAC INY-06-061, and surface MHC-I (upper panels) and total MHC-I were determined by FC and immunoblot, respectively. GAPDH levels are shown as loading control. (**D**) ERK5 PROTAC degrader does not alter MHC-I in cancer cells with high basal MHC-I membrane levels. ARK1, A549 and SW620 cells were treated and analyzed as in (**A)**. (**E**) ERK5 overexpression impairs MHC-I membrane expression in endometrial cancer cells. Ishikawa cells were co-transfected with vectors encoding for GST-tagged ERK5 and GFP, and MHC-I membrane expression was analyzed by FC within the GFP^+^ cell population. Left bar graphs show the percentage of MHC-I expressing cells and the MFI values. Immunoblots show levels of ERK5 and MHC-I. p21 protein expression was used as control for ERK5 activity, and Hsp90 as loading control. The numbers indicate relative MHC-I protein levels (mean ± SD from three independent experiments. Results represent mean ± SD from three independent experiments. ns, not significant; ** *p* < 0.005; **** *p* < 0.0001 as determined by one-way ANOVA followed by Bonferroni multiple comparisons test (**A-D**) or two-tailed Student’s *t*-test (**E**).

In addition, we also evaluated the effects of ERK5 and MEK5 inhibitors on MHC-I surface expression in 3D cultures of Ishikawa EC cells, a more complex model for drug testing. Flow cytometry analysis revealed that, as it happened for 2D cultures, treatment with the ERK5i or MEK5i or the ERK5 PROTAC increased the percentage of MHC-I+ cells and MFI values for MHC-I (**Fig. 3A-B**). Finally, we investigated whether ERK5 inhibition also affects MHC-I expression of cancer cells in vivo. To do so, we used tumor xenograft samples from a previous study, where the systemic administration 50 mg/kg JWG-071 in athymic nude mice impaired the growth of Ishikawa EC xenografts [16]. Immunohistochemical analysis of paraffin-embedded tumor sections revealed that systemic administration of the ERK5i resulted in higher number of tumor cells expressing membrane MHC-I (**Fig. 3C**).

**Figure 3.**
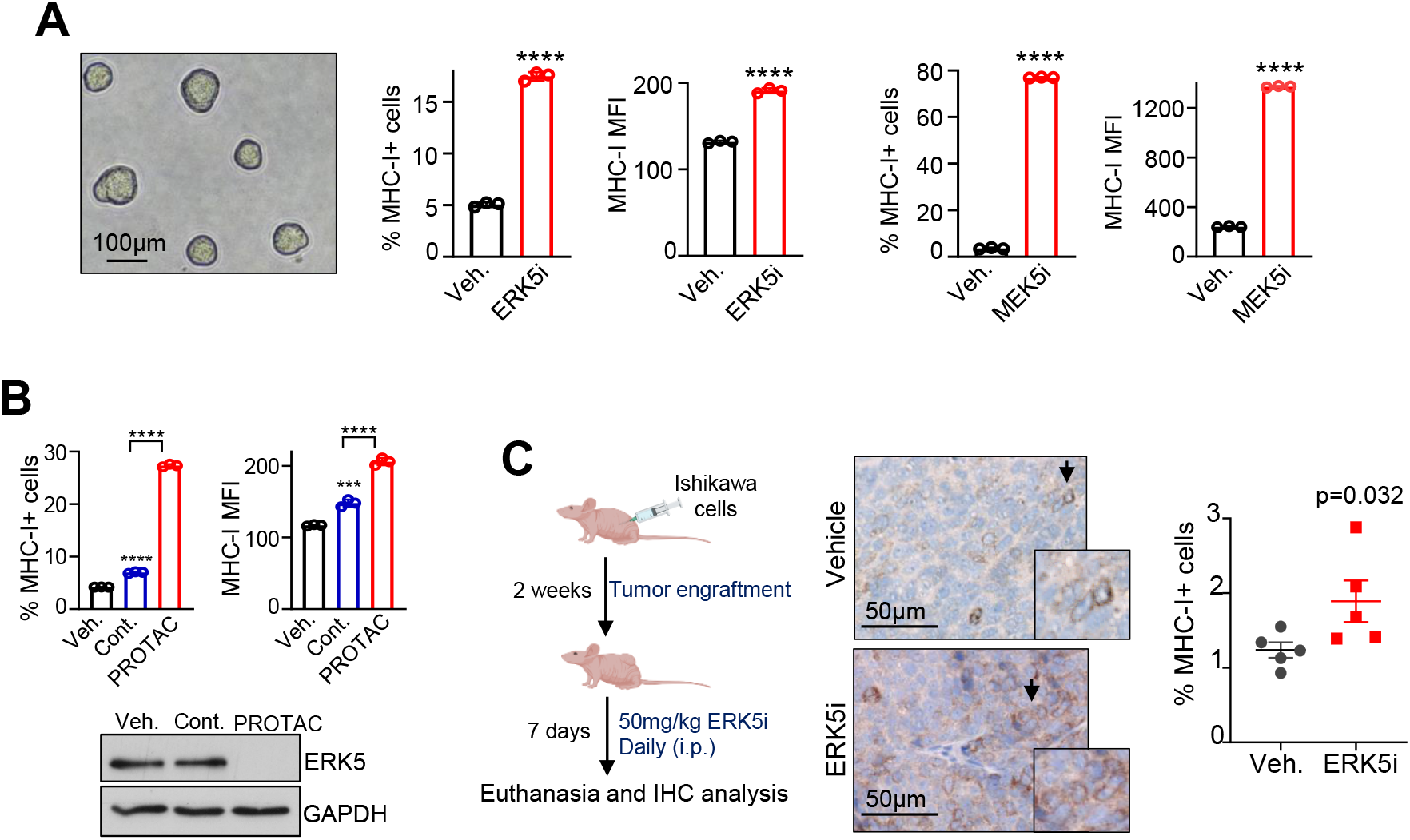
ERK5 inhibitor and PROTAC degrader enhance MHC-I membrane levels in 3D cultures of cancer cells and tumor xenografts. (**A, B**) 3D spheroid cultures of EC Ishikawa cells were treated for 24 h with either vehicle, 5μM JWG-071 or 5μM GW284543 (**A**), or with either vehicle, 300nM PROTAC negative control INY-06-089 (Cont.), or 300nM ERK5 PROTAC INY-06-061 (**B**). MHC-I surface expression was determined by FC analysis. (**A**) A representative image of untreated 3D cultures on day five is shown. Right histograms show the percentage of MHC-I expressing cells and MFI values. (**B**) Upper histograms show the percentage of MHC-I expressing cells and MFI values. Immunoblot shows ERK5 protein levels. **(A-B)** Data are the mean ± SD of a representative experiment from three independent experiments, each performed in triplicate. *** *p* < 0.001; **** *p* < 0.0001, as determined by two-tailed Student’s *t*-test (**A**) or one-way ANOVA followed by Bonferroni multiple comparisons test (**B**). (**B**) Athymic nude female mice were subcutaneously engrafted with Ishikawa cells. When tumors reached 80–100 mm^3^, mice were treated (i.p., daily) with either vehicle or 50 mg/kg JWG-071 for 7 days. Left histograms show the corresponding tumor volumes at day 7 (n=6 for each group) (**C**) Immunohistochemical analysis of the expression of surface MHC-I in tumors from nude mice engrafted with Ishikawa cells and treated either with vehicle or 50 mg/kg JWG-071 for 7 days. Scatter plot shows the percentage of cells expressing membrane MHC-I (*N*=5 mice per group, 5 fields per tumor; mean ± SD). Significance as determined by Mann-Whitney test.

Together, our results show that ERK5 inhibition upregulates MHC-I surface expression in cancer cells both in vitro and in vivo, supporting the potential of inhibitors of the ERK5 pathway to improve the immunogenic profile of tumor cells.

### ERK5 does not regulate transcription of genes encoding for MHC-I or for the antigen presentation pathway proteins

To investigate the mechanism by which ERK5 regulates MHC-I surface expression in cancer cells, we study the expression of genes encoding MHC-I molecules (*HLA-A, HLA-B, HLA-C* and *B2M*) and of the genes involved in the antigen presentation pathway (*TAP1* and *TAP2*) by RT-qPCR analysis. In Ishikawa EC cells, ERK5 or MEK5 inhibition did not alter the mRNA levels of any of these genes (**Fig. 4A**). Similarly, in IMR-32 neuroblastoma cells, gene expression remained mainly unchanged following treatment with ERK5 or MEK5 inhibitors, with the exception of *HLA-B* and *HLA-C*. In IMR-32 cells, ERK5 or MEK5 inhibitors enhanced *HLA-B* mRNA levels, while only the ERK5i led to increased *HLA-C* mRNA levels (**Fig. 4B**). As control, the ERK5i and MEK5i significantly reduced the mRNA levels of the ERK5 transcriptional target *KLF2* (**Fig. 4A-B**). Although ERK5i modestly enhanced the mRNA levels of *HLA-B* and *HLA*-*C* in IMR-32 cells, this is unlikely to have a significant impact on the levels of MHC-I expressed by these cells: our qPCR data found that IMR-32 cells mainly express *HLA-A* mRNA (Ct=21), whereas *HLA-B* and *HLA-C* mRNAs were barely detected (Ct: 36 and 38, respectively).

**Figure 4.**
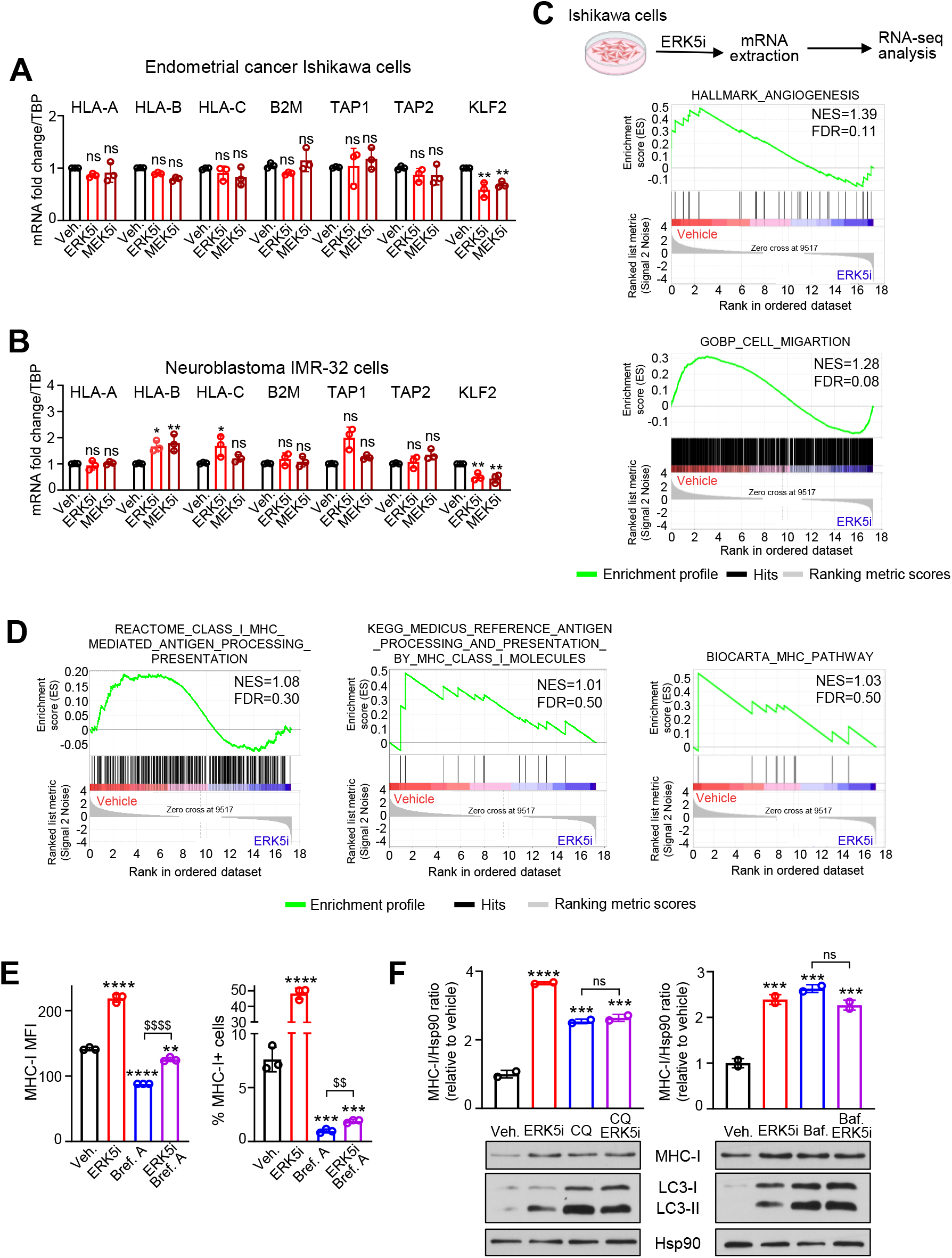
ERK5 does not regulate transcription of classical HLA-I genes or antigen-processing pathway. (**A, B**) ERK5/MEK5 inhibition does not alter mRNA levels of *HLA-A, HLA-B, HLA-C, B2M, TAP1* and *TAP2* genes. Ishikawa EC cells (**A**) or IMR-32 NBL cells (**B**) were treated for 24 hours with either vehicle, or 5μM JWG-071, or 5μM GW284543. Relative gene expression, normalized to *TBP* mRNA was assessed by RT-qPCR. *KLF2* relative expression was used as a control for ERK5 or MEK5 inhibition. (**C, D**) Effect of ERK5 inhibition on the transcriptional programs of Ishikawa EC cells. Cells were treated for 24 hours with 5μM JWG-071, followed by RNA-seq analysis. Panels show gene set enrichment analysis (GSEA) for the Hallmark “Angiogenesis” and the GoBP “Cell migration” (**C**), and for the Reactome “Class I MHC mediated antigen processing presentation” pathway, the KEGG “antigen processing and presentation by MHC Class I” pathway, and the Biocarta “MHC Pathway” (**D**). (**E**) ERK5 inhibition does not enhance Golgi-to plasma membrane transport of MHC-I. Ishikawa cells were treated for 24h with either vehicle, or 5μM JWG-071, or 250nM brefeldin A (Bref.A), or the combination of both, and MHC-I surface expression was determined by FC. Histograms show MHC-I MFI values and the percentages of cells expressing MHC-I. (**F**) Effect of ERK5 inhibition and of the lysosome inhibitors chloroquine (CQ) and bafilomycin A (Baf.), on MHC-I levels. Ishikawa cells were treated for 24h with either vehicle, or 5μM JWG-071, or 10μM chloroquine/150nM bafilomycin A), or the combination of both. Total MHC-I protein levels were determined by immunoblot analysis. Histograms show the quantification of MHC-I levels, referred to Hsp90. Autophagy marker LC3-I/II was used as a control for lysosome inhibitors, and Hsp90 as loading control. Data represent mean ± SD from three (**A, B, E**) or two (**F**) independent experiments. ns, not significant; * *p* < 0.05; **, $$ *p* < 0.005; *** *p* < 0.001; ****, $$$$ *p* < 0.0001, one-way ANOVA followed by Bonferroni multiple comparisons test.

To discard a potential role for ERK5 in the transcriptional regulation of other genes of the antigen presentation pathway, we also analyzed data from a RNA sequencing analysis of Ishikawa EC cells treated with JWG-071 [23]. Gene Set Enrichment Analysis (GSEA) revealed that ERK5 inhibition altered the transcriptional programs associated with angiogenesis and cell migration (**Fig. 4C**), two hallmarks regulated by ERK5 [8]. However, the ERK5i did not alter the transcriptional programs associated with antigen presentation pathway via MHC Class I (FDR > 0.25) (**Fig. 4D**). Overall, our results suggest that ERK5 may modulate the surface expression of MHC-I through posttranslational mechanisms, rather than altering the transcription of MHC-I genes or other components of the antigen presentation pathway.

Since transcriptional regulation does not account for the effects of ERK5 inhibition on MHC-I surface expression, we explored whether post-transcriptional processes such as trafficking to cell membrane or lysosomal degradation might be involved. First, we used the inhibitor of the Golgi-to-plasma membrane transport Brefeldin A [29]. As expected, Brefeldin A reduced the MHC-I membrane levels (**Fig. 4E**). However, ERK5 inhibitor still increased the MHC-I membrane expression in cells pre-treated Brefeldin A (**Fig. 4E**), suggesting that ERK5i does not enhance anterograde transport of MHC-I. We next examined the role of lysosomal degradation using the lysosomal inhibitors chloroquine and bafilomycin A. Both compounds increased total MHC-I protein levels and induced autophagy (**Fig. 4F**), as previously reported [30]. Of relevance, chloroquine or bafilomycin A did not further enhance the increase in total MHC-I protein levels induced by the ERK5i (**Fig. 4F**), indicating that ERK5 inhibition increases MHC-I expression by impairing its lysosomal degradation.

### ERK5 inhibition sensitizes cancer cells to the cytotoxic activity of tumor-specific CD8+ T cells by enhancing MHC-I surface expression

Next, we investigated whether the increased expression of surface MHC-I induced by ERK5i has functional relevance in enhancing antigen presentation and subsequent recognition and killing by cytotoxic CD8+ T cells. For this purpose, we next established an in vitro assay where cancer cells were co-cultured with human cancer cell-specific CD8+ T cells from three different donors. First, peripheral blood mononuclear cells (PBMCs) were incubated with tumor antigens derived from IMR-32 or Ishikawa cancer cell lines, along with GM-CSF and IL-4, to induce differentiation of monocytes to antigen-loaded dendritic cells (DCs). Then, T cell priming, activation and expansion was promoted with continuous culture for 15 days with IFN-γ, IL-2 and anti-CD3 antibody. After that, CD8+ T cells, predominantly tumor-specific T cells, were isolated by magnetic separation method. Finally, we co-cultured tumor-specific CD8+ T cells with cancer cells pre-treated with either vehicle or the ERK5i JWG-071 at different effector-to-target (E:T) ratios, and apoptosis in cancer cells was quantified by flow cytometry Annexin V/PI staining (**Fig. 5A**). As shown in **Fig. 5B, C and D**, when cultured alone, neither IMR-32 neuroblastoma nor Ishikawa endometrial cancer cells showed differences in apoptosis between vehicle and ERK5i treated groups. However, when co-cultured with tumor-specific CD8+ T cells for 18 hours, IRM-32 (**Fig. 5C)** and Ishikawa (**Fig. 5D and Supplementary Fig. 4**) cancer cells treated with the ERK5i showed increased proportion of apoptotic cells, compared with cancer cells not exposed to the ERK5 inhibitor. This increase was mainly due to the late apoptotic population, suggesting that ERK5 inhibition potentiates a rapid apoptosis induced by CD8+ T cells. Of relevance, the ERK5 inhibitor increased the levels of surface MHC-I in IRM-32 and Ishikawa cells co-cultured with CD8+ T cells **(Fig. 5E and F)**.

**Figure 5.**
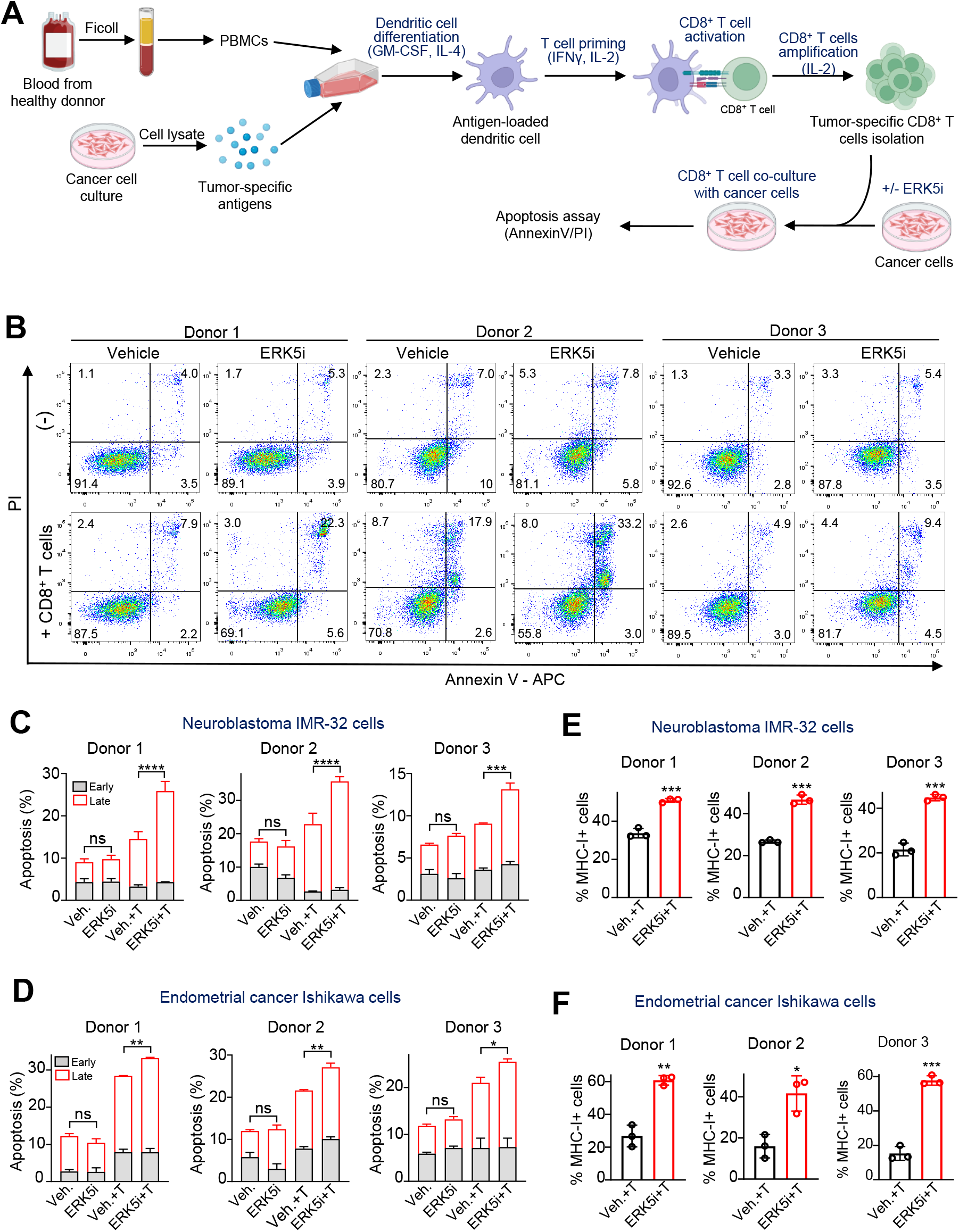
ERK5 inhibition enhances CD8+ T cell–mediated apoptosis of cancer cells. (**A**) Schematic representation of the protocol used to generate tumor-specific CD8+ T cells and their co-culture with human cancer cells. Tumor-specific antigens derived from IMR-32 or Ishikawa cancer cells were used to stimulate dendritic cells (DCs) derived from PBMCs of three healthy donors. Subsequently, T cells were activated and expanded. Finally, CD8+ T cells were isolated and co-cultured for 18 h with cancer cells pre-treated with either vehicle or JWG-071. The cytotoxic activity of CD8+ T cells in cancer cells was monitored by flow cytometry analysis of apoptosis (Annexin V/PI). (**B**) Representative dot plots of flow cytometry apoptosis assays for IMR-32 cells. (**C**) Bar graphs show the percentages of early (Annexin V+/PI-) and late (Annexin V+/PI+) apoptotic IMR-32 cells (n=3). Effector CD8+ T cells:target IMR-32 cells (E:T) ratios were 8:1 for donors 1 and 2, and 2:1 for donor 3. (**D**) Percentage of MHC-I+ IMR-32 cells treated with either vehicle or JWG-071 and co-cultured with tumor-specific CD8+ T cells from three different donors. (**E**) Percentages of early and late apoptotic cells (n=3) for flow cytometry apoptosis assays performed with Ishikawa cells. Effector CD8+ T cells:target Ishikawa cells (E:T) ratios were 16:1, 8:1 and 12:1, respectively. (**F**) Percentage of MHC-I+ Ishikawa cells treated with either vehicle or JWG-071 and co-cultured with tumor-specific CD8+ T cells from three different donors. Data represents mean ± SD from three independent experiments. **** *p* < 0.0001, *** *p* < 0.001, ** *p* < 0.005, * *p* < 0.05, determined by two-way ANOVA followed by Tukey multiple comparisons test.

Together, our results indicate that ERK5 pathway plays a role in regulating MHC-I surface expression, and that ERK5 inhibitors sensitize cancer cells to CD8+ T cell toxicity by enhancing T cell recognition by upregulating MHC-I membrane levels.

## DISCUSSION

Downregulation of surface MHC-I is a common mechanism used by cancer cells to escape recognition by CD8+ T cells, and is associated with poor prognosis and resistance to immune checkpoint blockade immunotherapy [3]. Understanding the mechanism underlying MHC-I deregulation is therefore crucial, and several anticancer strategies are currently investigated to restore MHC-I expression and enhance antigen presentation, such as treatment with interferons, epigenetic modulators or targeted inhibitors [31]. In this study, we identify the oncogenic MEK5-ERK5 signaling pathway as a repressor of MHC-I surface expression in a subset of human cancer cell lines with intrinsically low MHC-I and diverse mutational backgrounds, including NMYC-amplified neuroblastoma, PTEN/PI3KCA-mutant endometrioid cancer, and hormone-sensitive prostate cancer. Thus, ERK5 or MEK5 inhibition, as well as targeted degradation of ERK5, restored MHC-I surface and total protein levels in these cells. Moreover, systemic administration of an ERK5 inhibitor also increased MHC-I expression in tumor xenografts. In contrast, ERK5 over-expression, as it happens in some solid cancers [32], induced downregulation of surface MHC-I.

Mechanistically, our data suggest that EK5 regulates surface MHC-I levels through post-translational mechanisms. In PTEN/PI3KCA-mutant Ishikawa cells, inhibition of ERK5 or MEK5 increased membrane MHC-I expression without detectable changes in transcripts encoding MHC-I subunits or components of the antigen presentation machinery. Consistently, RNA-Seq analysis of the effect of the ERK5 inhibitor in these cells did not reveal significant alterations in transcriptional programs associated with the MHC-I antigen presentation pathway. In MYCN-amplified IMR-32 neuroblastoma cells, ERK5 and MEK5 inhibitors induced only a modest increase in HLA-B/C mRNA levels; however, given their very low basal expression, these changes are unlikely to account for the substantial restoration of MHC-I surface expression. In contrast, our preliminary data suggest that ERK5 impairs MHC-I lysosomal degradation, as two distinct lysosome inhibitors did not further enhance the increase in MHC-I levels induced by ERK5 inhibition. On the other hand, our preliminary data discard a role of ERK5 on the anterograde Golgi-to plasma membrane transport of MHC-I, since ERK5 inhibition increased surface MHC-I levels even in the presence of the anterograde transport inhibitor brefeldin A. Further work will be required to define the precise mechanism by which ERK5 regulates MHC-I proteostasis. In this regard, ERK5 has been reported to regulate the ubiquitylation and proteostasis of proteins such as TP53INP2 [23], MYC [28] or KLF2 [33]. On the other hand, lysosomal degradation of MHC-I is regulated by ubiquitination-dependent trafficking pathways mediated by several E3 ubiquitin ligases, including MARCH4/8 and the viralK3/K5 ligases [34]. Hence, a plausible mechanism is that ERK5 modulates MHC-I surface expression by regulating ubiquitin-dependent trafficking or the activity of specific E3 ligases.

Collectively, our findings point ERK5 as one of the first MAP kinases described to post-translationally regulate MHC-I in cancer cells, as most MAP kinases implicated in MHC-I regulation –such as MEK/ERK1/2 [20] or p38 [21]– act primarily at the transcriptional level, modulating the interferon/STAT1/NLRC5 signaling or epigenetic programs.

We found that ERK5 inhibition restored surface MHC-I across the well-established oncogenic contexts known to suppress antigen presentation and promote immune evasion: *MYCN* amplification [35], *PTEN/PIK3CA* mutations [36], and androgen receptor signaling [37]. These oncogenic pathways primarily regulate MHC-I levels at the transcriptional level. In this regard, although ERK5/MEK5 inhibitors altered c-Myc protein levels –but did not affect Akt or ERK1/2 activity-in the cell lines tested, their effect on MHC-I expression is post-translational.

Overall, our results highlight the potential broad applicability of ERK5 targeting to reverse MHC-I downregulation across diverse tumor types. Whether ERK5 inhibition potentiates MHC-I upregulation induced by inhibitors of c-Myc, Akt or ERK1/2 pathways deserves future investigation.

Our results contrast to a previous work showing that ERK5 silencing with retroviral vectors resulted in reduced MHC-I surface levels in human leukemic B cells by transcriptional mechanisms (impaired expression of β2M mRNA, [38]). We have no explanation for this discrepancy, which may reflect differences in tumor type (hematologic versus solid); however, our results are unlikely to reflect off-target effects of the MEK5 or ERK5 inhibitors used, since the exclusively selective ERK5 PROTAC compound INY-06-061 [22] also induced upregulation of MHC-I levels in our cell models. Moreover, our results are consistent with numerous works pointing ERK5 as an anticancer target [39].

Importantly, we show that ERK5 inhibition enhances CD8+ T cell-mediated antitumor immunity. Consistent with its role in regulating MHC-I expression, ERK5 inhibition sensitized endometrial cancer and neuroblastoma cells to CD8+ T cell-induced apoptosis. These results are in agreement with recent findings showing that ERK5 genetic depletion increases CD8+ T-cell infiltration in a syngeneic breast cancer model [40] and in a PTEN-deficient prostate cancer mouse model [41]. Although MHC-I upregulation increases cancer cell recognition by cytotoxic CD8+ T cells, it impairs the antitumor activity of Natural Killer (NK) cells mediated by the cytolytic pathway (granzyme b/perforin) that induces intrinsic apoptosis in cancer cells [42]. Interestingly, we recently showed that ERK5 inhibitors sensitized cancer cells to the antitumor activity of NK cells, promoting the extrinsic apoptosis induced by the death receptor agonists expressed by NKs [23], a pro-death pathway that is not affected by MHC-I surface expression in cancer cells [43]. Given that ERK5 overexpression or activation in cancer cells confers resistance to the death receptor agonists-induced apoptosis [23, 44], our results provide further evidence supporting an oncogenic role for the ERK5 pathway, blunting both NK cell- and CD8+ T cell-mediated immune surveillance.

Recent work showed that ERK5 silencing sensitizes triple-negative breast cancer to anti-PD1 immunotherapy in a syngenic model [40]). Since downregulation of MHC-I surface expression correlates with poor responses to immune checkpoint blockade, the results presented in this work support a rationale for the use of ERK5 inhibitors as a strategy to improve the efficacy of ICI-based immunotherapy.

## Conclusions

We used a panel of human cancer cell models representing three well-established major oncogenic contexts of reversible MHC-I downregulation (MYCN amplification, PTEN/PIK3CA mutations, and androgen receptor signaling) to identify ERK5 as a novel regulator of MHC-I in cancer cells. Pharmacologic inhibition of ERK5 restores MHC-I surface expression in cultured cancer cells and tumor xenografts with reversible low MHC-I levels, without affecting PD-L1, while functional assays with co-cultures of cancer cells with tumor-specific CD8+ T cells demonstrated that ERK5 blockade enhanced CD8+ T cell–mediated apoptosis of cancer cells. In contrast to other MAPKs RT-qPCR analysis showed that ERK5 pathway inhibition did not significantly modify transcription of classical HLA-I genes or antigen-processing machinery, but rather affected lysosomal degradation of MHC-I, supporting ERK5 as a new posttranslational regulator of MHC-I in cancer. Our results provide a rational for ERK5 inhibitors as a strategy to improve the efficacy of immune checkpoint blockade-based immunotherapy.

## List of abbreviations

DCs: Dendritic cells
DMEM: Dulbecco’s modified Eagle medium
FBS: Fetal Bovine Serum
FDR: False discovery rate
GM-CSF: Granulocyte-macrophage colony-stimulating factor
*GSEA:*: Gene set enrichment analysis
*HRP:*: Horseradish peroxidase
IFN: interferon
MAPK: Microtubule-associated MAP kinase
MEM: Modified Eagle medium
MHC: Major histocompatibility Complex
NEEA: Non-essential amino acid
NES: Normalized enrichment score
PBMC: Peripheral blood mononuclear cells
PBS: Phosphate buffered saline
PROTAC: Proteolysis-targeting chimera
RIPA: Radio-immunoprecipitation assay
RT-qPCR: Real time quantitative PCR

## Ethical approval and Consent to participate

Procedures with animals were conducted in compliance with the applicable laws, regulations, and guidelines of the Universitat Autònoma de Barcelona (UAB) and with the approval of the Ethics Committee on Animal Experiments from UAB (ref. CEEAH 3353) and VHIR (CEEAH 4342), according to Spanish official regulations. Buffy Coats were from the Banc de Sang i Teixits of Barcelona from volunteer blood donors, with the approval of the Committee of Ethics and Clinical Investigation of the Hospital Vall d’Hebron of Barcelona (PR(AG)449/2023).

## Consent to publish

Not applicable.

## Availability of data and materials

The RNA-seq datasets used and analyzed during the current study are available from the GEO database repository under accession code GSE239920.

## Authors’ contributions

M.V.-C. performed the experiments. M.V.-C. and J.M.L. conceived and designed the experiments, analyzed the data, prepare the figures and wrote the main manuscript text. J.M.L., resources and funding acquisition. Both authors read, critically revised, and approved.

## Acknowledgements

We thank Prof. Nathanael Gray (Stanford University, California) for generously providing the ERK5 degrader INY-06-061 and its negative control INY-06-089. We thank Dr. Lorena Usero for technical advice with flow cytometry and for work performed with CD8+T cells. M Viñas-Casas is a recipient of a FPU PhD fellowship funded by the Spanish Ministry of Science and Innovation (Ref. FPU2021-01999). The JM Lizcano research group was supported by grants from the Spanish Ministry of Science and Innovation (grant PID2019-107561RB-I00) and Spanish Ministry of Science, Innovation and Universities (grant PID2022-136391OB-100), and co-funded by the European Regional Development Fund (ERDF). We also thank to the UAT Service from VHIR.

## References

[1] N.M. Anderson, M.C. Simon, The tumor microenvironment, Curr Biol, 30 (2020) R921–R925.

[2] D. Hanahan, Hallmarks of Cancer: New Dimensions, Cancer Discov, 12 (2022) 31–46.

[3] F. Garrido, I. Algarra, A.M. Garcia-Lora, The escape of cancer from T lymphocytes: immunoselection of MHC class I loss variants harboring structural-irreversible “hard” lesions, Cancer Immunol Immunother, 59 (2010) 1601–1606.

[4] K. Dhatchinamoorthy, J.D. Colbert, K.L. Rock, Cancer Immune Evasion Through Loss of MHC Class I Antigen Presentation, Front Immunol, 12 (2021) 636568.

[5] B.H. Alsaafeen, B.R. Ali, E. Elkord, Resistance mechanisms to immune checkpoint inhibitors: updated insights, Mol Cancer, 24 (2025) 20.

[6] X. Wu, T. Li, R. Jiang, X. Yang, H. Guo, R. Yang, Targeting MHC-I molecules for cancer: function, mechanism, and therapeutic prospects, Mol Cancer, 22 (2023) 194.

[7] B.A. Drew, M.E. Burow, B.S. Beckman, MEK5/ERK5 pathway: the first fifteen years, Biochim Biophys Acta, 1825 (2012) 37–48.

[8] B. Stecca, E. Rovida, Impact of ERK5 on the Hallmarks of Cancer, Int J Mol Sci, 20 (2019).

[9] A.B. Bhatt, S. Patel, M.D. Matossian, D.A. Ucar, L. Miele, M.E. Burow, P.T. Flaherty, J.E. Cavanaugh, Molecular Mechanisms of Epithelial to Mesenchymal Transition Regulated by ERK5 Signaling, Biomolecules, 11 (2021).

[10] A. Sanchez-Fdez, M.F. Re-Louhau, P. Rodriguez-Nunez, D. Ludena, S. Matilla-Almazan, A. Pandiella, A. Esparis-Ogando, Clinical, genetic and pharmacological data support targeting the MEK5/ERK5 module in lung cancer, NPJ Precis Oncol, 5 (2021) 78.

[11] A. Sanchez-Fdez, S. Matilla-Almazan, S. Del Carmen, M. Abad, E. Arconada-Luque, J. Jimenez-Suarez, L.M. Chinchilla-Tabora, M.J. Ruiz-Hidalgo, R. Sanchez-Prieto, A. Pandiella, A. Esparis-Ogando, Etiopathogenic role of ERK5 signaling in sarcoma: prognostic and therapeutic implications, Exp Mol Med, 55 (2023) 1247–1257.

[12] J.W. Antoon, E.C. Martin, R. Lai, V.A. Salvo, Y. Tang, A.M. Nitzchke, S. Elliott, S.Y. Nam, W. Xiong, L.V. Rhodes, B. Collins-Burow, O. David, G. Wang, B. Shan, B.S. Beckman, K.P. Nephew, M.E. Burow, MEK5/ERK5 signaling suppresses estrogen receptor expression and promotes hormone-independent tumorigenesis, PLoS One, 8 (2013) e69291.

[13] K.G. Finegan, D. Perez-Madrigal, J.R. Hitchin, C.C. Davies, A.M. Jordan, C. Tournier, ERK5 is a critical mediator of inflammation-driven cancer, Cancer Res, 75 (2015) 742–753.

[14] A.K. Ramsay, S.R. McCracken, M. Soofi, J. Fleming, A.X. Yu, I. Ahmad, R. Morland, L. Machesky, C. Nixon, D.R. Edwards, R.K. Nuttall, M. Seywright, R. Marquez, E. Keller, H.Y. Leung, ERK5 signalling in prostate cancer promotes an invasive phenotype, Br J Cancer, 104 (2011) 664–672.

[15] M. Miranda, E. Rozali, K.K. Khanna, F. Al-Ejeh, MEK5-ERK5 pathway associates with poor survival of breast cancer patients after systemic treatments, Oncoscience, 2 (2015) 99–101.

[16] N. Dieguez-Martinez, S. Espinosa-Gil, G. Yoldi, E. Megias-Roda, I. Bolinaga-Ayala, M. Vinas-Casas, G. Gorgisen, I. Domingo-Orti, H. Perez-Montoyo, J.R. Bayascas, E. Colas, X. Dolcet, J.M. Lizcano, The ERK5/NF-kappaB signaling pathway targets endometrial cancer proliferation and survival, Cell Mol Life Sci, 79 (2022) 524.

[17] F. Tesser-Gamba, A.S. Petrilli, M.T. de Seixas Alves, R.J. Filho, Y. Juliano, S.R. Toledo, MAPK7 and MAP2K4 as prognostic markers in osteosarcoma, Hum Pathol, 43 (2012) 994–1002.

[18] E. Rovida, M.G. Di, I. Tusa, S. Cannito, C. Paternostro, N. Navari, E. Vivoli, X. Deng, N.S. Gray, A. Esparis-Ogando, E. David, A. Pandiella, S.P. Dello, M. Parola, F. Marra, The mitogen-activated protein kinase ERK5 regulates the development and growth of hepatocellular carcinoma, Gut, 64 (2015) 1454–1465.

[19] D.M. Pereira, C.M.P. Rodrigues, Targeted Avenues for Cancer Treatment: The MEK5-ERK5 Signaling Pathway, Trends Mol Med, 26 (2020) 394–407.

[20] E.J. Brea, C.Y. Oh, E. Manchado, S. Budhu, R.S. Gejman, G. Mo, P. Mondello, J.E. Han, C.A. Jarvis, D. Ulmert, Q. Xiang, A.Y. Chang, R.J. Garippa, T. Merghoub, J.D. Wolchok, N. Rosen, S.W. Lowe, D.A. Scheinberg, Kinase Regulation of Human MHC Class I Molecule Expression on Cancer Cells, Cancer Immunol Res, 4 (2016) 936–947.

[21] Z. Liu, F. Ning, Y. Cai, H. Sheng, R. Zheng, X. Yin, Z. Lu, L. Su, X. Chen, C. Zeng, H. Wang, L. Liu, The EGFR-P38 MAPK axis up-regulates PD-L1 through miR-675-5p and down-regulates HLA-ABC via hexokinase-2 in hepatocellular carcinoma cells, Cancer Commun (Lond), 41 (2021) 62–78.

[22] I. You, K.A. Donovan, N.M. Krupnick, A.S. Boghossian, M.G. Rees, M.M. Ronan, J.A. Roth, E.S. Fischer, E.S. Wang, N.S. Gray, Acute pharmacological degradation of ERK5 does not inhibit cellular immune response or proliferation, Cell Chem Biol, 29 (2022) 1630–1638 e1637.

[23] S. Espinosa-Gil, S. Ivanova, E. Alari-Pahissa, M. Denizli, B. Villafranca-Magdalena, M. Vinas-Casas, I. Bolinaga-Ayala, A. Gamez-Garcia, C. Faundez-Vidiella, E. Colas, M. Lopez-Botet, A. Zorzano, J.M. Lizcano, MAP kinase ERK5 modulates cancer cell sensitivity to extrinsic apoptosis induced by death-receptor agonists, Cell Death Dis, 14 (2023) 715.

[24] A. Gamez-Garcia, I. Bolinaga-Ayala, G. Yoldi, S. Espinosa-Gil, N. Dieguez-Martinez, E. Megias-Roda, P. Munoz-Guardiola, J.M. Lizcano, ERK5 Inhibition Induces Autophagy-Mediated Cancer Cell Death by Activating ER Stress, Front Cell Dev Biol, 9 (2021) 742049.

[25] T. Erazo, S. Espinosa-Gil, N. Dieguez-Martinez, N. Gomez, J.M. Lizcano, SUMOylation Is Required for ERK5 Nuclear Translocation and ERK5-Mediated Cancer Cell Proliferation, Int J Mol Sci, 21 (2020).

[26] X. Ma, Q. Wang, C. Sun, I. Agarwal, H. Wu, J. Chen, C. Zhao, G. Qi, Q. Teng, C. Yuan, S. Yan, J. Peng, R. Li, K. Song, Q. Zhang, B. Kong, Targeting TCF19 sensitizes MSI endometrial cancer to anti-PD-1 therapy by alleviating CD8(+) T cell exhaustion via TRIM14-IFN-beta axis, Cell Rep, 42 (2023) 112944.

[27] J. Wang, T. Erazo, F.M. Ferguson, D.L. Buckley, N. Gomez, P. Muñoz-Guardiola, N. Diéguez-Martínez, X. Deng, M. Hao, W. Massefski, O. Fedorov, N.K. Offei-Addo, P.M. Park, L. Dai, A. DiBona, K. Becht, N.D. Kim, M.R. McKeown, J.M. Roberts, J. Zhang, T. Sim, D.R. Alessi, J.E. Bradner, J.M. Lizcano, S.C. Blacklow, J. Qi, X. Xu, N.S. Gray, Structural and Atropisomeric Factors Governing the Selectivity of Pyrimido-benzodiazipinones as Inhibitors of Kinases and Bromodomains, ACS Chem Biol, (2018).

[28] A.V. Vaseva, D.R. Blake, T.S.K. Gilbert, S. Ng, G. Hostetter, S.H. Azam, I. Ozkan-Dagliyan, P. Gautam, K.L. Bryant, K.H. Pearce, L.E. Herring, H. Han, L.M. Graves, A.K. Witkiewicz, E.S. Knudsen, C.V. Pecot, N. Rashid, P.J. Houghton, K. Wennerberg, A.D. Cox, C.J. Der, KRAS Suppression-Induced Degradation of MYC Is Antagonized by a MEK5-ERK5 Compensatory Mechanism, Cancer Cell, 34 (2018) 807–822 e807.

[29] P. Munoz-Guardiola, J. Casas, E. Megias-Roda, S. Sole, H. Perez-Montoyo, M. Yeste-Velasco, T. Erazo, N. Dieguez-Martinez, S. Espinosa-Gil, C. Munoz-Pinedo, G. Yoldi, J.L. Abad, M.F. Segura, T. Moran, M. Romeo, J. Bosch-Barrera, A. Oaknin, J. Alfon, C. Domenech, G. Fabrias, G. Velasco, J.M. Lizcano, The anti-cancer drug ABTL0812 induces ER stress-mediated cytotoxic autophagy by increasing dihydroceramide levels in cancer cells, Autophagy., (2020) 1–18.

[30] E. Xanthopoulou, I. Lamprou, A.G. Mitrakas, G.D. Michos, C.E. Zois, A. Giatromanolaki, A.L. Harris, M.I. Koukourakis, Autophagy Blockage Up-Regulates HLA-Class-I Molecule Expression in Lung Cancer and Enhances Anti-PD-L1 Immunotherapy Efficacy, Cancers (Basel), 16 (2024).

[31] A.M. Cornel, I.L. Mimpen, S. Nierkens, MHC Class I Downregulation in Cancer: Underlying Mechanisms and Potential Targets for Cancer Immunotherapy, Cancers (Basel), 12 (2020).

[32] M. Monti, J. Celli, F. Missale, F. Cersosimo, M. Russo, E. Belloni, A. Di Matteo, S. Lonardi, W. Vermi, C. Ghigna, E. Giurisato, Clinical Significance and Regulation of ERK5 Expression and Function in Cancer, Cancers (Basel), 14 (2022).

[33] H.A. Brown, C.A.C. Williams, H. Zhou, D. Rios-Szwed, R. Fernandez-Alonso, S. Mansoor, L. McMulkin, R. Toth, R. Gourlay, J. Peltier, N. Dieguez-Martinez, M. Trost, J.M. Lizcano, M.P. Stavridis, G.M. Findlay, An ERK5-KLF2 signalling module regulates early embryonic gene expression and telomere rejuvenation in stem cells, Biochem J, 478 (2021) 4119–4136.

[34] J.A. Nathan, P.J. Lehner, The trafficking and regulation of membrane receptors by the RING-CH ubiquitin E3 ligases, Exp Cell Res, 315 (2009) 1593–1600.

[35] R. Bernards, S.K. Dessain, R.A. Weinberg, N-myc amplification causes down-modulation of MHC class I antigen expression in neuroblastoma, Cell, 47 (1986) 667–674.

[36] A.T. Parsa, J.S. Waldron, A. Panner, C.A. Crane, I.F. Parney, J.J. Barry, K.E. Cachola, J.C. Murray, T. Tihan, M.C. Jensen, P.S. Mischel, D. Stokoe, R.O. Pieper, Loss of tumor suppressor PTEN function increases B7-H1 expression and immunoresistance in glioma, Nat Med, 13 (2007) 84–88.

[37] L.N. Chesner, F. Polesso, J.N. Graff, J.E. Hawley, A.K. Smith, A. Lundberg, R. Das, T. Shenoy, M. Sjostrom, F. Zhao, Y.M. Hu, S. Linder, W.S. Chen, R.M. Hawkins, R. Shrestha, X. Zhu, A. Foye, H. Li, L.M. Kim, M. Bhalla, T. O’Loughlin, D. Kuzuoglu-Ozturk, J.T. Hua, M.L. Badura, S. Wilkinson, S.Y. Trostel, A.M. Bergman, D. Ruggero, C.G. Drake, A.G. Sowalsky, L. Fong, M.R. Cooperberg, W. Zwart, X. Guan, A. Ashworth, Z. Xia, D.A. Quigley, L.A. Gilbert, F.Y. Feng, A.E. Moran, Androgen Receptor Inhibition Increases MHC Class I Expression and Improves Immune Response in Prostate Cancer, Cancer Discov, 15 (2025) 481–494.

[38] S. Charni, B.G. de, M.G. Rathore, J.I. Aguilo, P.J. van den Elsen, D. Haouzi, R.A. Hipskind, J.A. Enriquez, M. Sanchez-Beato, J. Pardo, A. Anel, M. Villalba, Oxidative phosphorylation induces de novo expression of the MHC class I in tumor cells through the ERK5 pathway, J.Immunol., 185 (2010) 3498–3503.

[39] D.C. Miller, S.J. Harnor, M.P. Martin, R.A. Noble, S.R. Wedge, C. Cano, Modulation of ERK5 Activity as a Therapeutic Anti-Cancer Strategy, J Med Chem, 66 (2023) 4491–4502.

[40] C.O. Miranda M, Rozali E, Jeyaestiri S, Al-Muftah M and Al-Ejeh F, ERK5 Mediates Metastatic Colonization and Remodeling of The Tumor and Tumor-Immune Microenvironment and Immunotherapy Response, Fortune Journal of Health Sciences, 8 (2025).

[41] C.J. Loveridge, E.J. Mui, R. Patel, E.H. Tan, I. Ahmad, M. Welsh, J. Galbraith, A. Hedley, C. Nixon, K. Blyth, O. Sansom, H.Y. Leung, Increased T-cell Infiltration Elicited by Erk5 Deletion in a Pten-Deficient Mouse Model of Prostate Carcinogenesis, Cancer Res, 77 (2017) 3158–3168.

[42] I. Prager, C. Watzl, Mechanisms of natural killer cell-mediated cellular cytotoxicity, J Leukoc Biol, 105 (2019) 1319–1329.

[43] A. Ramirez-Labrada, C. Pesini, L. Santiago, S. Hidalgo, A. Calvo-Perez, C. Onate, A. Andres-Tovar, M. Garzon-Tituana, I. Uranga-Murillo, M.A. Arias, E.M. Galvez, J. Pardo, All About (NK Cell-Mediated) Death in Two Acts and an Unexpected Encore: Initiation, Execution and Activation of Adaptive Immunity, Front Immunol, 13 (2022) 896228.

[44] J. Borges, A. Pandiella, A. Esparis-Ogando, Erk5 nuclear location is independent on dual phosphorylation, and favours resistance to TRAIL-induced apoptosis, Cell Signal, 19 (2007) 1473–1487.

