## Supplementary Figures and Tables for "MAPK ERK5 Is a Novel Regulator of MHC-I in Cancer Cells"

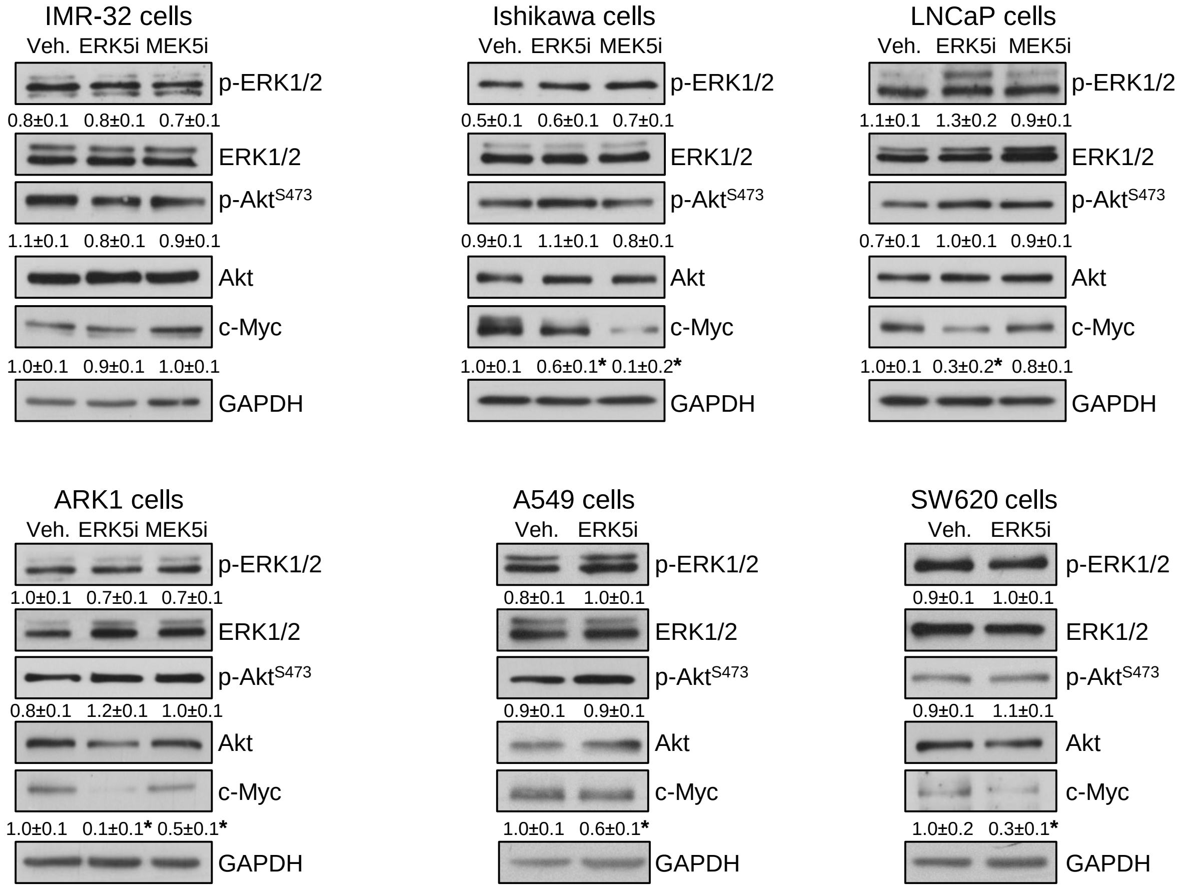


**Supplementary Figure 1.** Effect of ERK5 or MEK5 inhibitors in the ERK1/2, Akt and c-Myc pathways in cancer cells. Cells were incubated for 24h with 5 μM JWG-071 (ERK5i) or 5 μM GW2 (MEK5i). Representative immunoblots showing the levels of the indicated proteins and phosphoproteins. Numbers are the mean ± SD of at least two independent experiments. Levels of c-Myc are referred to vehicle. Levels of phosphorylated ERK1/2 or Akt are referred to total ERK1/2 or Akt protein levels, respectively *, $p<0.05$.

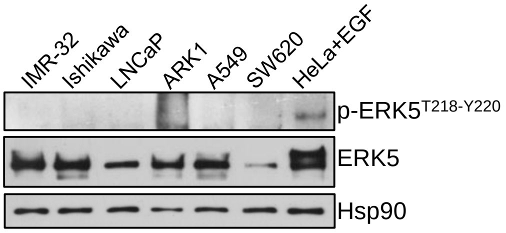


**Supplementary Figure 2.** Immunoblot analysis of total and phosphorylated ERK5 protein in lysates from human cancer cell lines. A lysate from HeLa cells treated with 50 ng/ml EGF for 20 min is shown as a control of phosphorylated ERK5.

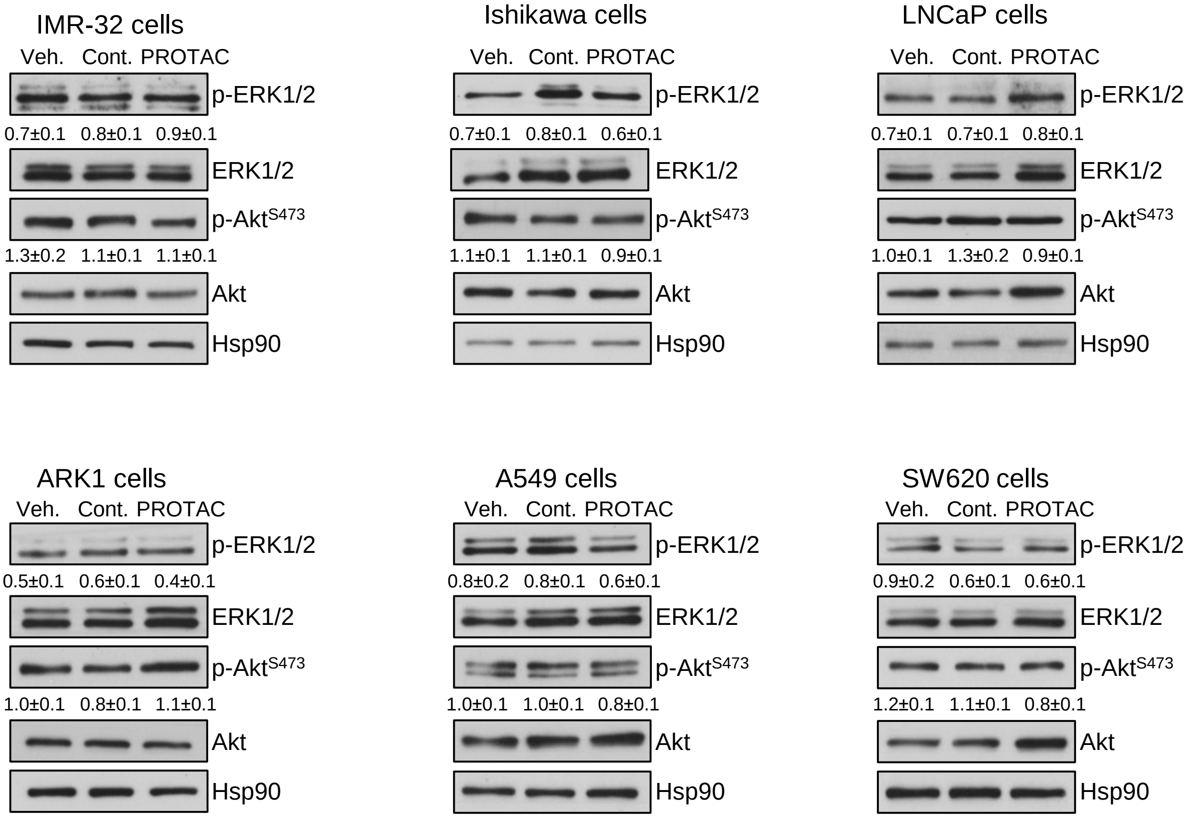


**Supplementary Figure 3.** Effect of the ERK5 PROTAC INY-06-061 or the control compound (Cont., INY-06-089) in the ERK1/2 and Akt pathways in cancer cells. Cells were incubated for 24h with 300 nM of each compound. Representative immunoblots showing the levels of the indicated proteins and phosphoproteins. Numbers are the mean ± SD of at least two independent experiments. Levels of phosphorylated ERK1/2 or Akt are referred to total ERK1/2 or Akt protein levels, respectively *, $p<0.05$.

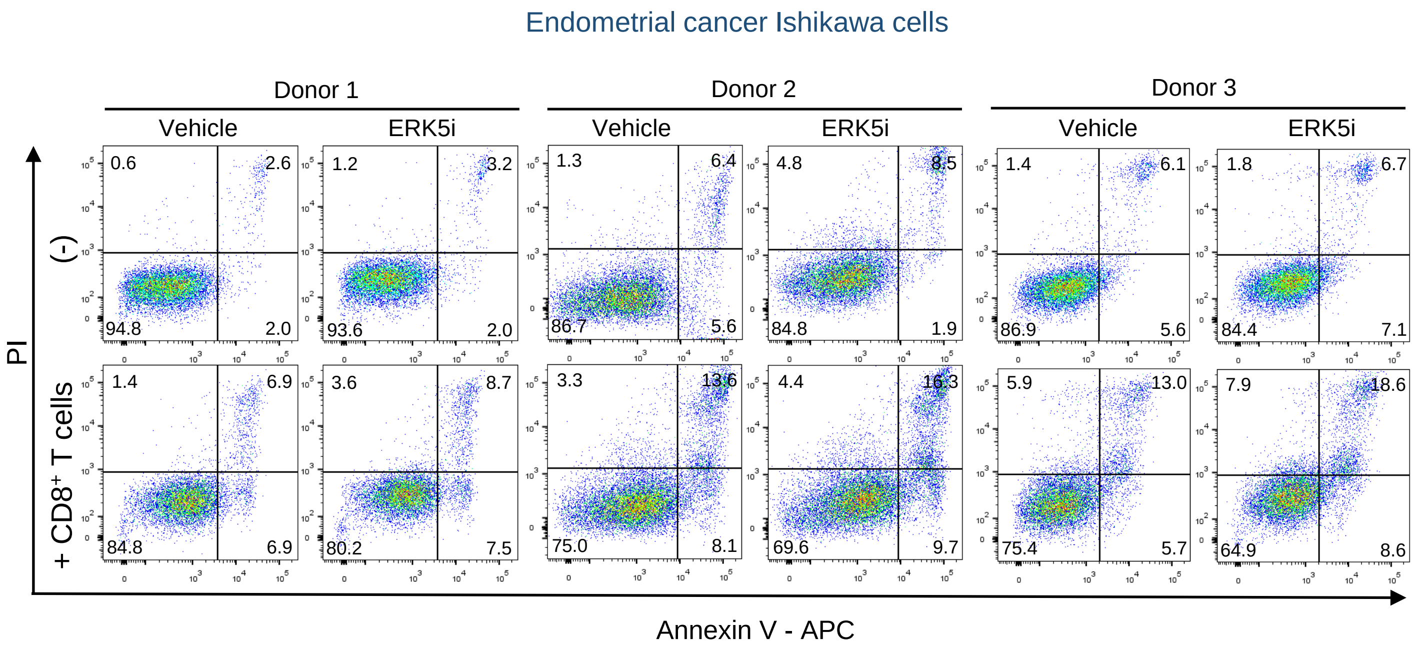


**Supplementary Figure 4**. ERK5 inhibition enhances CD8+ T cell–mediated apoptosis of human endometrial cancer cells. Representative dot plots of flow cytometry apoptosis assays for Ishikawa cells.

**Supplementary Table 1.** Primary and secondary antibodies used in immunoblot analyses.

| Antibody | Source | Reference | Dilution | Host |
|---|---|---|---|---|
| MHC-I | Cell Signaling | 35923 | 1:2000 | Rabbit |
| p21 | Millipore | 05-655 | 1:1000 | Mouse |
| p-ERK5 | Cell Signaling | 3371 | 1:1000 | Mouse |
| ERK5 | Cell Signaling | 3372 | 1:1000 | Rabbit |
| LC3I/II | Cell Signaling | 12741 | 1:2000 | Rabbit |
| p-ERK1/2 | Cell Signaling | 4376 | 1:1000 | Rabbit |
| ERK1/2 | Cell Signaling | 4695 | 1:8000 | Rabbit |
| p-Akt (S473) | Cell Signaling | 9271 | 1:1000 | Rabbit |
| Akt | Cell Signaling | 9272 | 1:5000 | Rabbit |
| c-Myc | Cell Signaling | 9402 | 1:5000 | Rabbit |
| Hsp90 beta | Invitrogen | PA3-012 | 1:16000 | Rabbit |
| GAPDH | Invitrogen | AM4300 | 1:100000 | Mouse |
| Goat anti-Rabbit IgG (H+L) Secondary Ab, HRP | Pierce | 31460 | 1:6000 | Goat |
| Goat anti-Mouse IgG (H+L) Secondary Ab, HRP | Pierce | 31430 | 1:6000 | Goat |

**Supplementary Table 2.** Amplification primers used in RT-qPCR assays.

| Gene name | Primer Forward (5'-3') | Primer Reverse (5'-3') |
|---|---|---|
| *HLA-A* | AGATACACCTGCCATGTGCAGC | GATCACAGCTCCAAGGAGAACC |
| *HLA-B* | CTGCTGTGATGTGTAGGAGGAAG | GCTGTGAGAGACACATCAGAGC |
| *HLA-C* | GGAGACACAGAAGTACAAGCGC | ACATCCTCTGGAGGGTGTGAGA |
| *B2M* | CCACTGAAAAAGATGAGTATGCCT | CCAATCCAAATGCGGCATCTTCA |
| *TAP1* | GCAGTCAACTCCTGGACCACTA | CAAGGTTCCCACTGCTTACAGC |
| *TAP2* | ATGCCCTTCACAATAGCAGCGG | CCAAAACTGCGAACGGTCTGCA |
| *KLF2* | CCAAGAGTTCGCATCTGAAGGC | CCGTGTGCTTTCGGTAGTGGC |
| *CDKN1A* | AGGTGGACCTGGAGACTCTCAG | TCCTCTTGGAGAAGATCAGCCG |
| *TBP* | GAATATAATCCCAAGCGGTTTG | ACTTCACATCACAGCTCCCC |

**Supplementary Table 3.** Cell lines used in this study and their major driver genetic alterations.

| **Cell line** | **Tumor type** | **Major driver alterations** |
|---|---|---|
| IMR-32 | Neuroblastoma | *MYCN* amplification; 1p loss; 17q gain |
| Ishikawa | Endometrioid cancer | *PTEN* loss; *TP53* mutation, *PIK3R1* mutation |
| LNCaP | Prostate cancer | *AR* T877A; *PTEN* loss; *PIK3CA* mutation |
| ARK1 | Serous endometrial cancer | *PIK3CA* mutation, *HER2* overexpression |
| A549 | Non-small cell lung cancer | *KRAS* G12S; *STK11* loss; *KEAP1* mutation |
| SW620 | Colorectal cancer | *KRAS* G12V; *APC* truncation; *TP53* mutation; *SMAD4* mutation |
